# HDAC6 regulates expression of the oncogenic driver EWSR1-FLI1 through the *EWSR1* promoter in Ewing sarcoma

**DOI:** 10.1101/2021.01.04.425179

**Authors:** Daniel J. García-Domínguez, Nabil Hajji, Sara Sánchez-Molina, Elisabet Figuerola-Bou, Rocío M. de Pablos, Ana M. Espinosa-Oliva, Eduardo Andrés-León, Laura Carmen Terrón-Camero, Rocío Flores-Campos, Guillem Pascual-Pasto, María José Robles, Ángel M. Carcaboso, Jaume Mora, Enrique de Álava, Lourdes Hontecillas-Prieto

## Abstract

Ewing sarcoma (EWS) is an aggressive developmental sarcoma driven by a fusion gene, EWSR1-FLI1. However, little is known about the regulation of EWSR1-FLI1 chimeric fusion gene expression. Here, we demonstrate that active nuclear HDAC6 in EWS modulates acetylation status of specificity protein 1 (SP1), consequently regulating SP1/P300 activator complex binding to EWSR1 and EWSR1-FLI1 promoters. Selective inhibition of HDAC6 impairs binding of activator complex SP1/P300, thereby inducing EWSR1-FLI1 downregulation and significantly reducing its oncogenic functions. In addition, sensitivity of EWS cell lines to HDAC6 inhibition is higher than other tumor or non-tumor cell lines. Overexpression of HDAC6 in primary EWS tumor clinical samples correlates with a poor prognosis. Notably, a combination treatment of a selective HDAC6 inhibitor and doxorubicin (standard of care in EWS) dramatically inhibits tumor growth in two EWS murine xenograft models. These results could lead to suitable and promising therapeutic alternatives for EWS patients.

## INTRODUCTION

Ewing sarcoma (EWS) is a highly aggressive and poorly differentiated rare tumor that affects children and young adults (Grunewald *et al*, 2018). EWS tumors usually contain a t(11;22)(q24;q12) chromosomal translocation involving the *EWSR1* gene (Ewing sarcoma breakpoint region 1) and ETS transcription factors. This reciprocal translocation generates a fusion oncogene, with *EWSR1-FLI1* being the most common (Delattre *et al*, 1992; Grunewald *et al.*, 2018). Continuous expression of EWSR1-FLI1 in a permissive cellular context is essential both for preserving the malignant phenotype in EWS cells and for their survival, and its inhibition induces apoptosis (Gierisch *et al*, 2019; Herrero-Martin *et al*, 2009; Kinsey *et al*, 2006; Martinez-Lage *et al*, 2020; Prieur *et al*, 2004; Smith *et al*, 2006). As EWS tumors have a remarkably stable genome at time of diagnosis (Crompton *et al*, 2014), the role of EWSR1-FLI1 in the EWS epigenome regulation has been deeply explored. Even though little is known about the molecular mechanisms responsible for regulating the gene expression of the chimeric fusion, researchers have looked for strategies to target the EWSR1-FLI1 activity that is critical for EWS tumorigenesis (Kovar, 2010; Tanaka *et al*, 1997). This chimeric fusion protein acts as an aberrant transcription factor, deregulating the expression of numerous target genes, which are then involved in EWS disease progression (Grunewald *et al.*, 2018; Riggi *et al*, 2014; Sankar *et al*, 2013b). EWSR1-FLI1 can recruit different proteins that participate in its transcription regulator function, such as lysine-specific histone demethylase 1A (LSD1) and histone deacetylases (HDACs) (Sankar *et al*, 2013a). In addition, the chimeric protein regulates other essential mechanisms, thereby compromising the epigenetic homeostasis; these include modulating enhancers (Riggi *et al.*, 2014; Sanchez-Molina *et al*, 2020), regulating exon splicing (Selvanathan *et al*, 2015), reprogramming metabolism(Tanner *et al*, 2017), and deregulating histone acetylation (both by repressing histone acetyltransferases [HAT] and by enhancing HDAC activities)(Sakimura *et al*, 2005).

The involvement of HDACs in cancer led them to be considered as optimal drug targets, leading to the development of several HDAC inhibitors for use in the clinic (Sun *et al*, 2018). Recently, we have shown that the pan-HDAC inhibitor SAHA inhibits EWSR1-FLI1 expression (Garcia-Dominguez *et al*, 2018). However, it remains to be determined which HDAC(s) are involved specifically in the regulation of EWSR1-FLI1 expression. Usually, the treatment with non-selective HDAC inhibitors in patients is an imprecise mechanism associated with increased toxicity (Hontecillas-Prieto *et al*, 2020). Hence, the identification of specific HDAC(s) involved in the regulation of EWSR1-FLI1, and the use of selective inhibitors, may improve efficacy and avoid toxicity of treatments, and enhance our understanding about the regulatory mechanisms of EWSR1-FLI1.

HDAC6, a class II HDAC, has not been specifically described in EWS. HDAC6 activity had been fundamentally observed in the cytoplasm (Lee *et al*, 2008), where α-tubulin and HSP90α have been defined as its substrates (Seidel *et al*, 2015). These proteins are involved in protein trafficking, protein degradation, cell morphology, and migration, and deregulation of HDAC6 activity is associated with a variety of diseases, including cancer (Lee *et al.*, 2008; Seidel *et al.*, 2015). Although HDAC6 localization is partially nuclear (Liu *et al*, 2012), it is not clear if it plays a functional role in gene regulation via the regulation of histone acetylation. In fact, several studies revealed a role of HDAC6 in regulating the activity of transcription factor modulators (Wang *et al*, 2009; Yang *et al*, 2015). In our recent study, we identified that HDAC6 specifically regulates histone acetylation at a specific residue, H4K12, in several types of cancer, including EWS (García-Domínguez, 2020). We determined that an increased acetylation level in H4K12 following selective HDAC6 inhibition induces a major chromatin relaxation and sensitizes cancer cells to DNA damage. From a clinical point of view, HDAC6 is overexpressed in many tumor types, and its overexpression is associated with poor prognosis, drug resistance, and cancer cell proliferation and migration (Kanno *et al*, 2012; Wang *et al*, 2016; Yang *et al.*, 2015). Further, knocking down HDAC6 in mice has no effects on viability or fertility (Zhang *et al*, 2008). Overall, these results make HDAC6 a potential target for EWS treatments.

As epigenetics plays a crucial role in EWS oncogenesis, we performed a screen assay of 43 epigenetic drugs in seven EWS cell lines to determine whether any of them prevents the oncogenic activities of EWSR1-FLI1. Our screen revealed that EWS cell lines were mostly sensitive to the specific HDAC6 inhibitor BML-281, which inhibited cell proliferation and subsequently induced apoptotic cell death. This provided our rationale to investigate in-depth the role and mechanism of HDAC6 in EWS cells *in vivo* and *in vitro*. This study presents for the first time clear evidence that HDAC6 specifically regulates the expression of *EWSR1-FLI1* by modulating the binding of the SP1/P300 complex to the *EWSR1-FLI1* promoter region. Our results, together with EWS patient data, provide strong evidence for an important role of HDAC6 in EWS, and suggest the use of selective HDAC6 inhibitors as a novel therapeutic approach to treat this type of sarcoma.

## RESULTS

### EWS cell lines are highly sensitive to selective inhibition of HDAC6

We and others have previously shown that HDAC inhibitors hinder EWS proliferation both *in vitro* and *in vivo*, using preclinical EWS models (Garcia-Dominguez *et al.*, 2018; Sakimura *et al.*, 2005). To investigate the key HDACs that control EWS sensitivity to HDAC inhibitors, we performed a screening assay for 43 compounds that affect HDAC activity (Screen-Well® Epigenetics Library BML-2836, Enzo). We used seven EWS cell lines to estimate the proliferation inhibition capacity of these compounds, and we calculated a median IC50 value for each drug and cell line (fig. S1A). The analysis revealed that only trichostatin A (a pan-HDAC inhibitor) and BML-281 (a selective HDAC6 inhibitor) had an IC50 value lower than 1.5 μM for all seven EWS cell lines (fig. S1B). Due to the well-documented toxicity and side effects of pan-HDACs inhibitors, we selected BML-281 rather than trichostatin A as a potent promising HDAC inhibitor for further analysis. We next expanded the list panel to thirteen EWS cell lines and validated the results of the compound screen for cells treated with BML-281. The median IC50 values in EWS cell lines (IC50 = 0.5465 μM) were significantly lower than those obtained for other tumor entity cell lines (IC50 = 0.8518 μM) and the non-tumor cell line hMSC (IC50 = 3.113 μM) (Fig. 1, A and B).

**Fig. 1.**
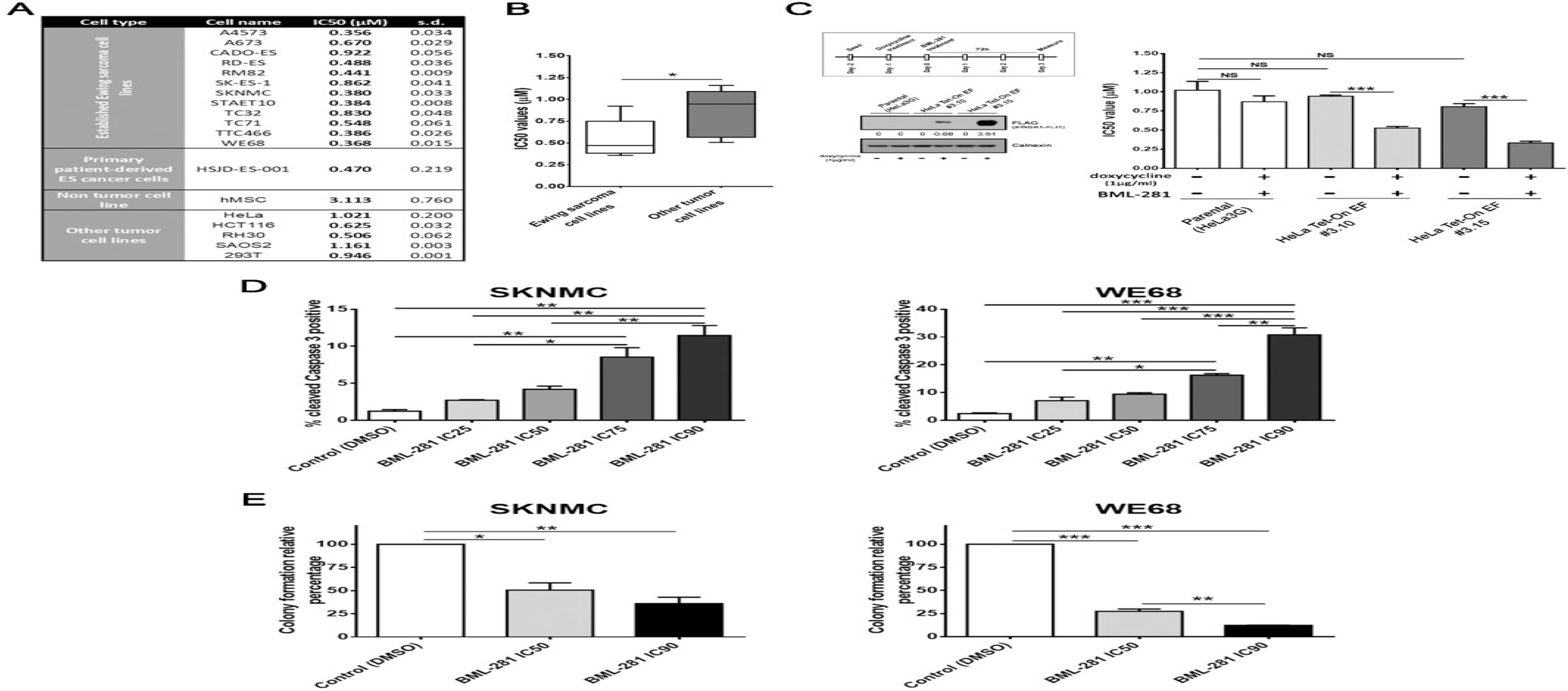
EWS cell lines showed a high sensitivity to selective HDAC6 inhibition. (**A**) Cell viability assessment by 50% inhibitory concentration (IC50) of 13 EWS cell lines, one non-tumor cell line, and 5 non-EWS cancer cell lines, after 72-hour exposure to BML-281. (**B**) Statistical comparison of the average sensitivity to IC50 HDAC6 inhibition between the two cell lines groups. (**C**) A schematic representation of the performed protocol for *EWSR1-FLI1* induction (upper left panel), and confirmation of the EWSR1-FLI1 ectopic expression protein by Western blotting using HeLa cells treated with doxycycline (lower left panel). Numbers below blots represent densitometric quantification of bands, normalized to endogenous bands. Statistical comparison of sensitivity to HDAC6 inhibition with ectopic EWSR1-FLI1 protein expression and no expression in two clones with inducible chimeric protein-expression system in HeLa cells (right panel). (**D**) Cleaved-caspase 3 level assessment in SKNMC and WE68 cells treated with BML-281 in dose-dependent manner for 48 hours and analyzed by flow cytometry. (**E**) Effects of HDAC6 inhibition on the anchorage-independent growth capacity of the EWS cell lines (SKNMC and WE68) by soft-agar colony formation assay. All values show mean ± s.d. of three biological independent replicates. Statistical tests were done using significant analysis of variance, Tukey’s post hoc test; \**P* < 0.05; \*\**P* < 0.01; \*\*\**P* < 0.001.

We next analyzed whether EWS cell sensitivity to the selective HDAC6 inhibitor BML-281 correlated with the level of HDAC6 protein expression. We demonstrated that there were no statistical differences in the levels of HDAC6 protein expression in the EWS cell lines as compared to other tumor cell lines (fig. S1, C and D). Therefore, our results showed that the sensitivity of EWS cells to selective inhibition of HDAC6 via BML-281 did not involve changes in its expression (fig. S1E). The significant sensitivity to HDAC6 selective inhibition in the EWS cell lines could be due to a role of the fusion protein as a regulator of the epigenetic landscape, as it is well known that EWSR1-FLI1 modulates HAT and HDAC activities and subsequently affects their targets (Sakimura *et al.*, 2005). To investigate whether the fusion protein expression “per se” sensitizes EWS cells to HDAC6 inhibition, we induced the chimeric protein expression in an ectopic model using HeLa cells. Notably, we previously demonstrated that EWSR1-FLI1 expression in HeLa cells mirrors its function in EWS (Garcia-Dominguez *et al*, 2020). The EWSR1-FLI1 expression in the two clones selected (#3.10 and #3.15) increased the sensitivity with respect to a non-induced system and reduced the IC50-BML-281 values (Fig. 1C). In contrast, the IC50 value in parental HeLa cells was not modified by doxycycline treatment. In sum, these results suggest that EWSR1-FLI1 expression in the doxycycline-independent HeLa model increased the sensitivity of HeLa cells to BML-281 treatment.

As BML-281 impairs proliferation in EWS cell lines, we analyzed its effects on physiological functions, such as apoptotic cell death and clonogenicity. For this, we determined the selective inhibition of HDAC6 effects on cleaved caspase 3 induction (apoptosis executor) in EWS cells after a 48-hour exposure to BML-281 by flow cytometry. We detected a significant induction of cleaved caspase 3 in different EWS cell lines, mainly at medium (IC50, IC75) and high (IC90) concentrations of BML-281, after a 48-hour exposure as compared to the control (Fig. 1D). We then investigated the residual effects of selective HDAC6 inhibition in a clonogenic assay (which measures the ability of EWS cells to form 3D-colonies in a soft agar medium) after a 24-hour exposure to medium or high doses of BML-281. At the IC50 and IC90 concentrations, colony formation was significantly reduced to 50.59% and 36.01%, respectively, in SKNMC cells, and to 27.42% and 12.22%, respectively, in WE68 cells, as compared to non-treated cells (Fig. 1E). Our results indicated that the selective inhibition of HDAC6 by BML-281 induced apoptosis via caspase 3 activation and reduced the capacity of cells for tumorigenesis *in vitro*. These results suggest that HDAC6 could be a potential target for EWS treatments.

### Nuclear HDAC6 activity inhibits EWSR1-FLI1 expression in EWS cell lines

As EWS cell lines are highly sensitive to HDAC6 inhibition, we analyzed the role of HDAC6 by measuring its activity. We first evaluated HDAC6 expression in response to BML-281 at different time points. The early time points of 4, 12, and 24 hours of BML-281 exposure did not affect protein or mRNA HDAC6 expression levels in either SKNMC or WE68 cells, as shown by immunoblotting, RT-qPCR, and RNA-seq results (Fig. 2A, fig. S2A, and Table 1). However, HDCA6 expression levels were reduced after a long-term (48 hour) exposure to BML-281 (Fig. 2A). Although the selective inhibitor BML-281 did not decrease HDAC6 expression with short-term exposure, HDAC6 activity was impaired. In both EWS cell lines, α-tubulin acetylation was strongly induced after HDAC6 inhibition in EWS cells lines after short-term treatment, and maintained over time (fig. S2B).

**Fig. 2.**
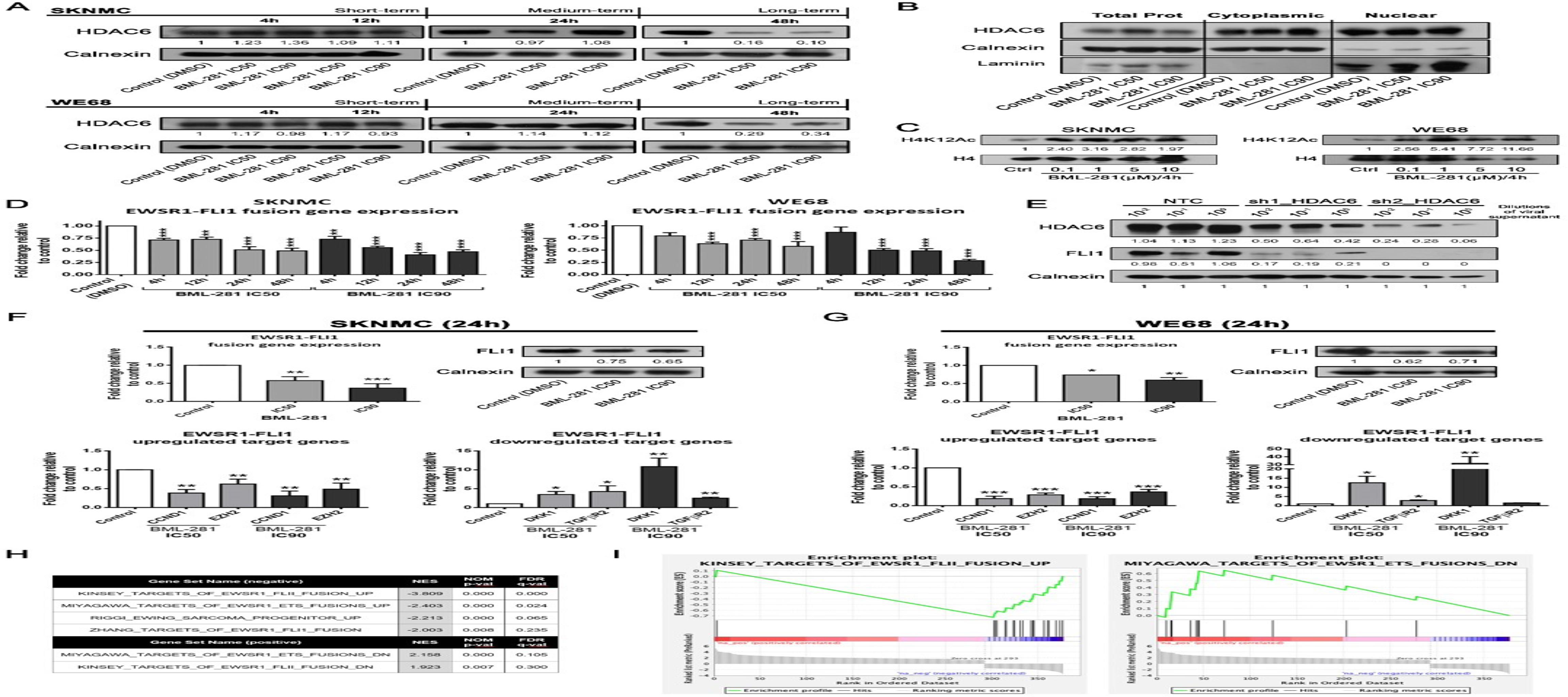
Inhibition of HDAC6 activity induces EWSR1-FLI1 downexpression and reduced its oncogenic activity. (**A**) Time course experiments (short-, medium-, and long-term) of HDAC6 protein expression was evaluated by immunoblotting using extracts from SKNMC and WE68 EWS cell lines treated with IC50 or IC90 concentrations of BML-281. (**B**) Subcellular fractionation and immunoblotting evaluation of HDAC6 localization in both cytoplasm and nucleus fractions. (**C**) Dose-dependent evaluation of H4K12 acetylation level in SKNMC and WE68 cell lines exposed to increasing concentrations of BML-281 (0.1, 1, 5, and 10 μM) for 4 hours. (**D**) RT-qPCR analysis of *EWSR1-FLI1* mRNA level in SKNMC and WE68 EWS cell lines treated with BML-281 (IC50 and IC90) in time-course experiments. (**E**) Immunoblot assessment of HDAC6 and EWSR1-FLI1 protein expression levels after HDAC6 depletion by two shRNA constructs. (**F** and **G**) RT-qPCR and immunoblot assessment of mRNA and protein expression levels of EWSR1-FLI1 (F and G, upper panels), and EWSR1-FLI1 regulating target genes (F and G, lower panels) after 24 hour of BML-281 treatment at IC50 and IC90 concentrations in the SKNMC or WE68 cell line, respectively. (**H**) GSEA C2_MSigDB analysis showed the overlap between genes that were significantly repressed or induced, by at least 1.5-fold, by BML-281 in EWS cell lines at EWSR1-FLI1 target genes signatures. (**I**) Enrichment plots with the best enrichment based on the |NES| (normalized enrichment score) are shown. Numbers below blots represent densitometric quantification of bands, normalized to endogenous bands and referred to their respective controls (DMSO band from the same time point). All values show mean ± s.d. of three biological independent replicates. Statistical tests were done using significant analysis of variance, Tukey’s post hoc test; \**P* < 0.05; \*\**P* < 0.01; \*\*\**P* < 0.001.

Although HDAC6 activity was described to be predominantly cytoplasmic, we confirmed the presence of both nuclear and cytoplasmic HDAC6 in EWS cell lines (Fig. 2B) with no changes in HDAC6 expression levels after a 24-hour BML-281 treatment in either cytoplasm or nucleus. We also explored the nuclear HDAC6 activity in EWS by evaluating the deacetylation of H4K12ac, which we have previously shown to be the specific histone-residue target (García-Domínguez, 2020). Selective HDAC6 inhibition strongly induced H4K12 residue acetylation in EWS cell lines (Fig. 2C). Further, we confirmed the nuclear HDAC6 role by using the enrichment_GO analysis (Table 2). These bioinformatics analyses interestingly revealed that early HDAC6 inhibition (at 4 hours) induced change in expression of genes involved in histone and chromatin modifications and in regulation of DNA binding of transcription factors. In addition, we observed that nuclear genes enrichment was maintained over 24 hours (Table 2). These data support the nuclear role of HDAC6 in EWS cells.

Due to nuclear localization of HDAC6 and its role in EWS cell lines, we further evaluated if HDAC6’s nuclear activity affects the fusion protein EWSR1-FLI1, which is considered the main driving force of EWS disease. Our results showed significant downregulation of *EWSR1-FLI1* mRNA at early time points after BML-281 treatment, starting at 4 hours in SKNMC, and 12 hours in WE68 cells (Fig. 2D). We also identified that BML-281 reduced the fusion protein expression level at 12 hours in both EWS cell lines, and this reduction was maintained over time (fig. S2C). To verify the specificity of BML-281 in reducing the levels of EWSR1-FLI1, HDAC6 expression was depleted by two shRNA constructs (Fig. 2E). Indeed, targeted HDAC6 downregulation specifically reduced the aberrant transcription factor expression, confirming that HDAC6 positively regulated EWSR1-FLI1.

We next checked the downstream effects of HDAC6’s regulation of EWSR1-FLI1 on expression of known EWSR1-FLI1 target genes (namely, *CCND1*, *EZH2*, *DKK1*, and *TGFβR2*). In parallel, we evaluated the downregulation of EWSR1-FLI1 (at both mRNA and protein levels). Genes that are canonically activated by upregulated EWSR1-FLI1 (*CCND1* and *EZH2*) were downregulated by HDAC6 inhibition, while those that are classically repressed (*DKK1* and *TGFβR2*) were upregulated, in both EWS cell lines at medium and high concentrations of BML-281 treatment (Fig. 2, F and G, and fig. S2, D and E).

To determine the relevance of EWSR1-FLI1 depletion by HDAC6 inhibition, we expanded our analysis to include genes that were significantly modulated by BML-281 in the SKNMC and WE68 cell lines (Table 1). After conducting a pre-ranked study with GSEA software, we compared our list of differentially expressed genes (with *P* < 0.05 and FC > |1.5|) against the C2_MSigDB signature database. We observed a statistically significant enrichment in EWS signatures (Kinsey, Miyagawa, Riggi, or Zhang), with negative normalized enrichment scores (NES) for mostly upregulated genes (UP gene sets), and positive NES for downregulated genes (DN gene sets), in our treated EWS cell lines (Fig. 2, H and I, and Table 3). These data revealed an inverse response to de novo expression of EWSR1-FLI1 in a previously described ectopic model (Garcia-Dominguez *et al.*, 2020). GSEA of rank-ordered gene analyses confirmed that the gene signature induced by HDAC6 inhibition recapitulated the molecular features, and therefore the biology, of the specific chimeric fusion depletion in EWS cell lines.

Altogether, we showed that *EWSR1-FLI1* downregulation is not due to a global transcription inhibition by BML-281 treatment; rather, our results indicated that EWSR1-FLI1 is downregulated by HDAC6 inhibition. In fact, we demonstrated that EWSR1-FLI1 downstream target genes were affected in response to EWSR1-FLI1 downregulation.

### HDAC6 regulates *EWSR1-FLI1* expression through SP1/P300 binding to the *EWSR1* promoter

Increased transcription levels of mirRNA-145 and Let-7 family members reduces the transcription levels of *EWSR1-FLI1* (Ban *et al*, 2011; Keskin *et al*, 2020). Therefore, we next explored whether EWSR1-FLI1 regulation by HDAC6 inhibition is due to the transcription modulation of mirRNA-145 and Let-7 family members. The RT-qPCR results showed that miR-145 expression levels were not significantly affected by BML-281, neither at increased concentrations nor at different exposure times (Fig. 3A). Therefore, we ruled out miR-145 as a possible intermediate between HDAC6 and EWSR1-FLI1. An analogous assay was performed for the remaining *Let-7* gene family (*let-7a*, *let-7b*, and *let-7c*). We observed an increase of *Let-7* family gene expression at 24 hours, followed by reduction at 48 hours, in the SKNMC cell line. However, we only observed a significant upregulation of the *Let-7* gene family at 48 hours in the WE68 cell line (Fig. 3B). Similar regulation of *Let-7* gene family expression was induced by IC50 and IC90 levels of BML-281 in both cell lines (Fig. 3B). These results suggested that Let-7 could contribute, at least in part, to EWSR1-FLI1 downregulation induced by HDAC6 selective inhibition in EWS cell lines. However, upregulation of *Let-7* gene family expression was consistent in EWS only after long-term HDAC6 inhibition and cannot explain EWSR1-FLI1 inhibition after short-term exposure to HDAC6 inhibition.

**Fig. 3.**
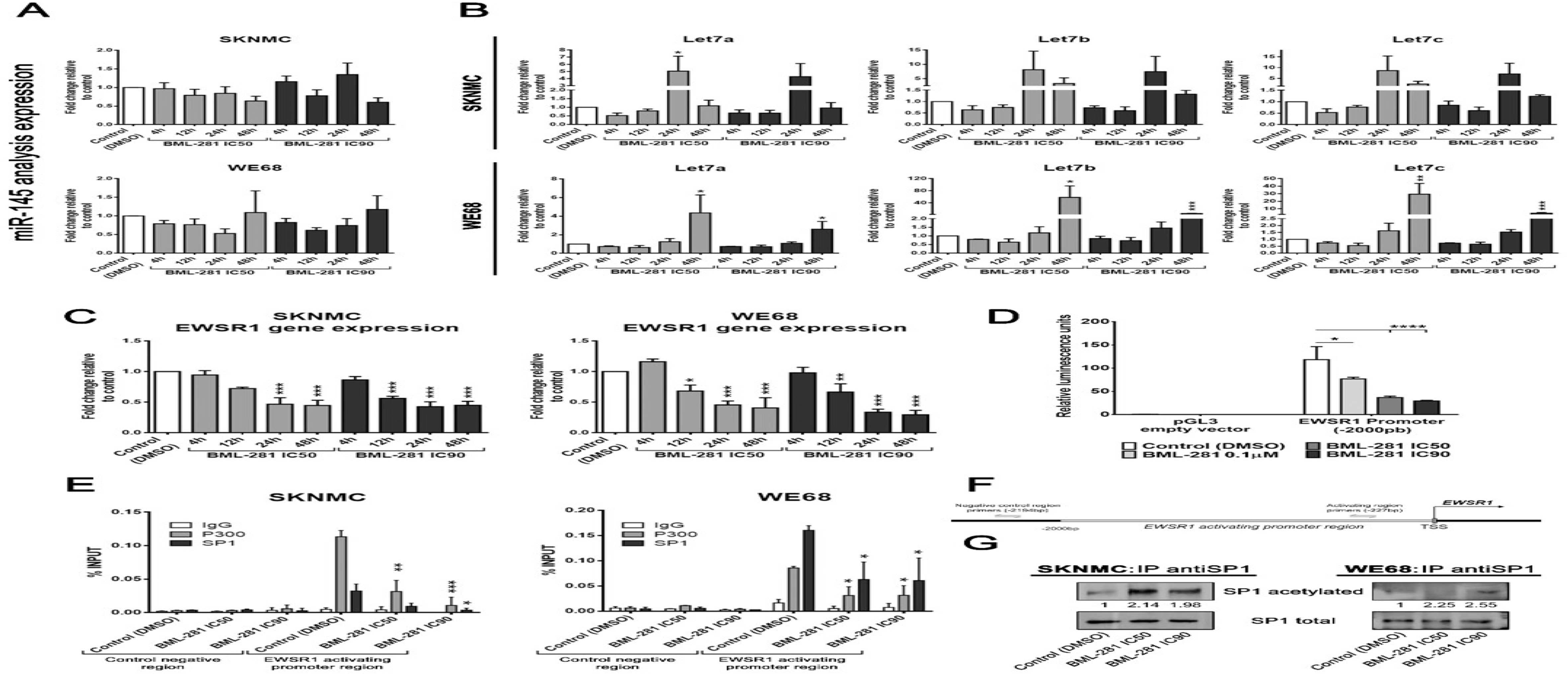
HDAC6 regulates *EWSR1-FLI1* and endogenous *EWSR1* expression through SP1/P300 binding to their promoters. (**A**) RT-qPCR analysis of miR-145 levels in SKNMC and WE68 EWS cell lines treated with BML-281. (**B**) RT-qPCR analysis of the expression levels of *Let-7* family members in SKNMC and WE68 EWS cell lines treated with BML-281 treatment. (**C**) RT-qPCR analysis of *EWSR1* mRNA expression of the non-translocated gene in SKNMC and WE68 EWS cell lines treated with BML-281. (**D**) Evaluation of *EWSR1* promoter region-dependent transcriptional activation with luciferase reporter constructs containing the region of 2000 bp upstream of the *EWSR1* transcription start site (TSS), in a dose-dependent BML-281-treatment of SKNMC cells for 4 hours. (**E** and **F**) ChIP and qPCR analysis of SP1 and/or P300 binding levels at EWSR1 activating promoter region in SKNMC and WE68 EWS cell lines treated with BML-281. % Input indicates enrichment ratio of immunoprecipitated samples relative to input. The IgG antibody was used as a control for unspecific binding in ChIP assays. Binding sites of the primers used are shown schematically (F). (**G**) SP1 pull-down and evaluation of lysine acetylated levels against total SP1 protein expression in both SKNMC and WE68 cell lines treated for 24 hours with increasing concentrations of BML-281 (IC50 and IC90). Numbers below blots represent densitometric quantification of bands, normalized to endogenous bands (total SP1) and referred to their respective controls (DMSO). All values show mean ± s.d. of three biological independent replicates. Statistical tests were done using significant analysis of variance, Tukey’s post hoc test; \**P* < 0.05; \*\**P* < 0.01; \*\*\**P* < 0.001.

To investigate regulation that occurs in the short-term, we used the ectopic EWSR1-FLI1 expression model in HeLa cells. In this model, the *EWSR1* promoter, characteristic of the fusion gene, was replaced by an inducible, doxycycline-dependent promoter. Surprisingly, BML-281 treatment induced fusion protein overexpression (fig. S3A). This response rules out posttranscriptional regulation as a principal *EWSR1-FLI1* inhibition mechanism by BML-281, and suggests that the gene fusion promoter is a potential candidate. Based on this indirect evidence, we further evaluated the role of the *EWSR1* promoter as a principal element for HDAC6-mediated *EWSR1-FLI1* inhibition. Consistent with the above results, we analyzed the endogenous (non-translocated) *EWSR1* expression using specific primers (that were previously described) (Garcia-Dominguez *et al.*, 2018). We observed a statistically significant downregulation after 12 hours or 24 hours of BML-281 treatment in WE68 and SKNMC cells, respectively (Fig. 3C); this is in agreement with the inhibition of *EWSR1-FLI1* expression shown by both RT-qPCR and RNA-seq (Fig. 2D and Table 1). To confirm *EWSR1*/*EWSR1-FLI1* promoter regulation by HDAC6, we designed a luciferase-reporter construct whose promoter comprised of 2000 bp upstream of the *EWSR1* transcription start site. The results showed a significant inhibition of luciferase expression in a dose-dependent manner (Fig. 3D). Altogether, our results demonstrated that regulation of expression of both *EWSR1* and *EWSR1-FLI1* by HDAC6 inhibition was mediated through the promoter, since both gene promoters (endogenous and translocated) have a homologous regulatory DNA sequence.

To better understand how HDAC6 regulates the *EWSR1*/fusion gene promoter, we conducted an *in silico* prediction of possible intermediate elements that could participate in this regulation. We found that the p300 protein is situated between HDAC6 and EWSR1 (fig. S3B). Previous studies demonstrated that EWSR1-FLI1 recruits p300 for activating gene expression (Riggi *et al.*, 2014). We therefore analyzed the physical interactions between HDAC6 and the fusion protein to exclude a possible positive feedback loop. Our results showed that there was no direct protein–protein binding (fig. S3C), suggesting the existence of protein(s) that facilitate p300 binding to the fusion protein promoter. Notably, it has been demonstrated that SP1 can recruits p300 to activate gene transcription (Azahri *et al*, 2012; Formisano *et al*, 2015; Hung *et al*, 2006). Further, SP1 is a *EWSR1-FLI1* positive regulator (Giorgi *et al*, 2015). In addition, the *in silico* analysis demonstrated that HDAC6, EWSR1, and SP1 are interconnected through p300 (fig. S3D). First, we evaluated *SP1* expression by both RNA-seq and RT-qPCR. Our results showed a slight downregulation of *SP1* after BML-281 treatment (Table 1 and fig. S3E) that was however not enough to explain the inhibition of fusion gene expression. To test whether HDAC6 regulates SP1 binding to EWSR1 and EWSR1-FLI1 promoters independently of the modulation of SP1 expression level, we analyzed binding of SP1 as well as of its coactivator P300 at activating region of *EWSR1* promoter (–2000 bp – 0; *EWSR1* transcription start site (Moller *et al*, 2009)) by chromatin immunoprecipitation (ChIP). We showed that SP1 and p300 binding was impaired by HDAC6 inhibition in both EWS cell lines analyzed (Fig. 3, E and F). Acetylation of lysines in SP1 is associated with its reduced DNA-binding capacity (Waby *et al*, 2010). Therefore, we analyzed the alteration of the lysine acetylation status of SP1 under BML-281 treatment through a pull-down assay. We demonstrated increased total lysine acetylation of SP1 by HDAC6 inhibition in both EWS cell lines (Fig. 3F). Together, these results support that HDAC6 regulates, at least partially, the expression of *EWSR1*/*EWSR1-FLI1* through SP1-P300 complex binding to its promoters.

### High HDAC6 expression in EWS tumor samples was associated with poor prognosis

To evaluate the impact of HDAC6 expression levels on prognosis of EWS patients, we first evaluated their expression levels in human EWS tumor samples by immunohistochemistry (IHC). Specific HDAC6 staining was found in both the nucleus and cytoplasm of EWS tumor cells (fig. S4A). Due to the different expression levels of HDAC6, the Kaplan–Meier analysis was conducted, grouping patient samples with negative/weak intensity versus moderate/strong intensity. EWS patients with moderate/strong intensity of HDAC6 had worse overall survival (OS) and disease-free survival rates as compared to the negative/weak intensity group (Fig. 4A). An additional *in silico* study using R2 genomics analysis and visualization platform (http://r2.amc.nl) confirmed these results and showed that EWS patients with high *HDAC6* expression had markedly lower OS and disease-free survival rates than those with low levels of *HDAC6* (fig. S4B). These findings indicated that high HDAC6 expression levels are associated with poor outcome for EWS patients.

**Fig. 4.**
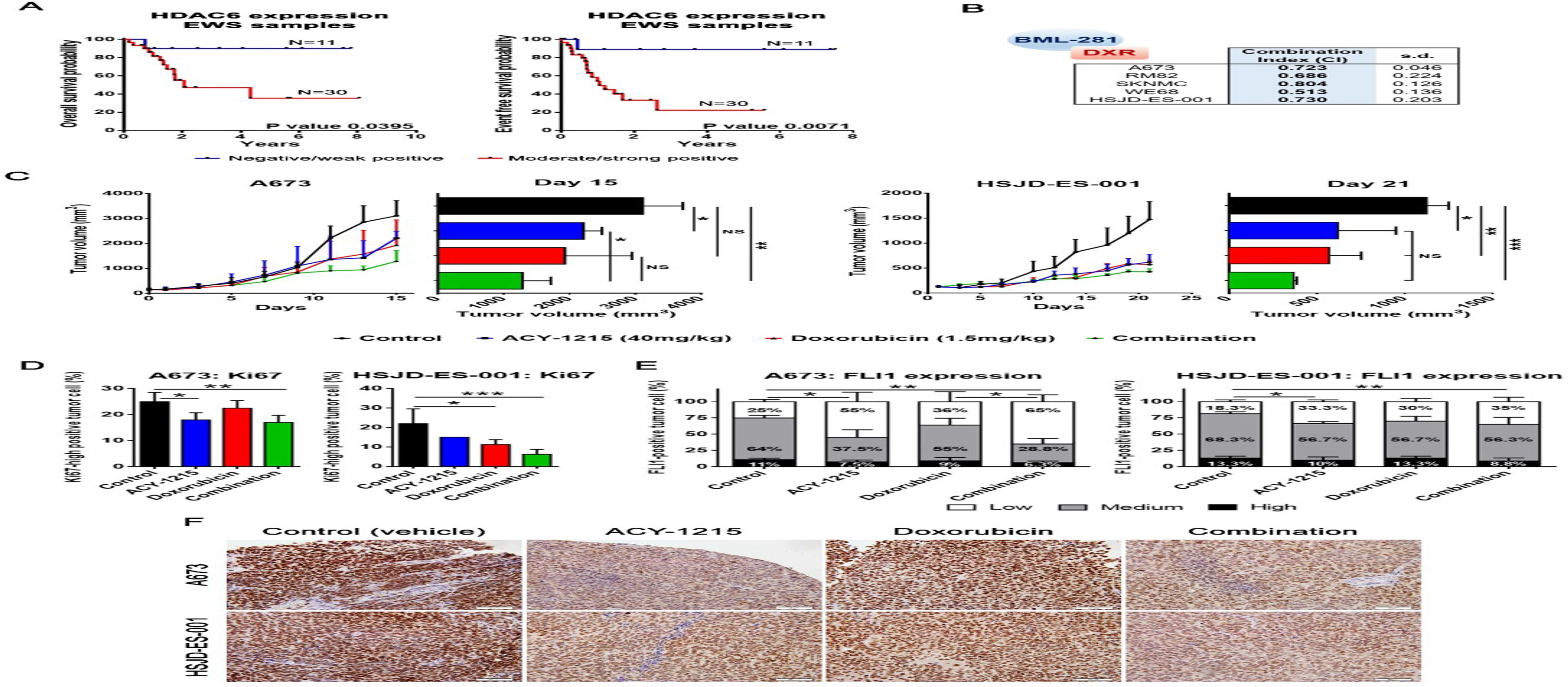
HDAC6 overexpression correlates with poor prognosis in EWS patients, and *in vivo* combined treatment reduces EWS tumor growth. (**A**) HDAC6 Kaplan– Meier plot with OS and DFS, according to the transcript. IHC expression in primary EWS tumor samples showed statistically significant differences between low- and high-HDAC6-expressing tumors. (**B**) Combination index (CI) of BML-281 and doxorubicin on proliferation inhibition in five EWS cell lines. (**C**) Tumor growth was monitored in A673 and HSJD-ES-001 xenografts models after single-agent or combination therapies. Tumor volumes (mm^3^) were measured after 15 days (A673) or 21 days (HSJD-ES-001) of treatment. (**D**) Quantification of Ki67-high positively-labelled nuclei percentage after 15 or 21 days of treatment. (**E**) Anatomopathological quantification of FLI1 expression after 15 or 21 days of treatment, with three levels of expression threshold set: low, medium, and high. (**F**) Immunohistochemical staining of FLI1 in xenograft tumor samples treated with ACY-1215 and doxorubicin, alone or in combination (40× magnifications). Statistical tests were done using significant analysis of variance, Tukey’s post hoc test; \**P* < 0.05; \*\**P* < 0.01; \*\*\**P* < 0.001.

**Fig. 5.**
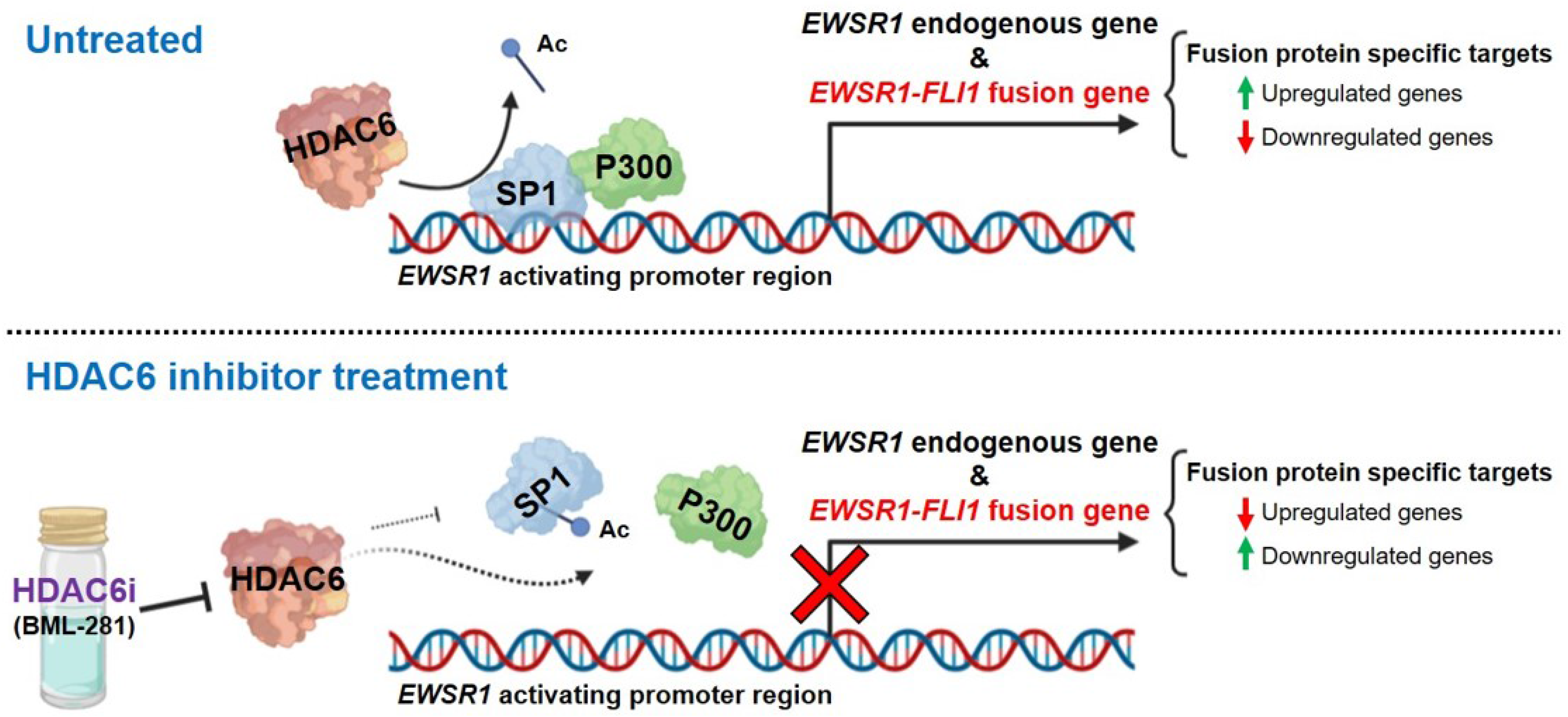
Schematic representation of regulation of *EWSR1* and *EWSR1-FLI1* transcription by SP1 and p300 binding complexes in both homologous promoter regions. HDAC6 modulates acetylation of SP1 as well as its binding to the *EWSR1*/*EWSR1-FLI1* activating promoter regions. In turn, HDAC6 increases SP1/p300 complex binding by deacetylating SP1. HDAC6 inhibition allows dissociation of the SP1/p300 complex by increased acetylation of SP1, with a subsequent decrease in both the expression of EWS fusion gene transcription and its protein activity, leading to modulation of a specific EWS target genes. Figure created with BioRender.com.

### HDAC6 inhibition impairs EWS tumor growth *in vivo*

Given the prognostic significance of HDAC6 overexpression in EWS patients, we hypothesize that its inhibition could be a promising new alternative as a clinical treatment. Therefore, we first conducted an *in vitro* experiment to evaluate the effect of the combination between doxorubicin (a cytotoxic drug used in standard EWS patient chemotherapy treatment) and HDAC6 inhibition (using BML-281). A strong synergistic effect (0.1 < combination index (CI) ≤ 0.7) was observed in the proliferation inhibition in both established and primary EWS cell line cultures from EWS patient-derived xenografts (PDX) (Fig. 4B).

Based on the strongly synergistic effects observed *in vitro* assays, we explored the therapeutic potential of HDAC6 inhibition (alone or in combination) *in vivo*. The A673 and HSJD-ES-001 cell lines were injected subcutaneously into CB-17 scid mice. Six days after injection, mice were divided randomly into four groups: control (vehicle), ACY-1215 (50 mg/kg), doxorubicin (1.5 mg/kg), or both ACY-1215 and doxorubicin (fig. S4C). ACY-1215 (also known as ricolinostat) is a potent and selective HDAC6 inhibitor (Seidel *et al.*, 2015; Wang *et al*, 2018); we used it in the *in vivo* experiment based on its antitumor effects and efficacy, both alone and in combination with various conventional treatments for cancer (Amengual *et al*, 2015; Cao *et al*, 2018; Santo *et al*, 2012) and in several clinical trials (Seidel *et al.*, 2015; Wang *et al.*, 2018). Treatment was well tolerated in mice, with no weight loss observed (fig. S4D), and histopathological evaluation did not reveal overall tissue alteration (fig. S4E). After three weeks of therapy, the tumor volume was significantly reduced to similar degrees in both cell lines for the monotherapy treatment groups (ACY-1215 or doxorubicin) as compared to the control group. Notably, the combined treatment in both EWS tumor models led to a greater reduction in tumor size than either monotherapy alone (Fig. 4C).

After mice necropsy, all tumors were isolated and processed. The histopathological examination using hematoxylin and eosin (H&E) staining reveled no morphological alterations of EWS cells in any analyzed samples (fig. S4F). However, the percentage of high Ki67-positive labelled cell populations in the combination treatment group was significantly lower than that of the control group, and both monotherapy groups had levels between the two (Fig. 4D). Finally, we evaluated if the *in vivo* assay replicated the EWSR1-FLI1 depletion induced by HDAC6 inhibition *in vitro*. An anti-FLI1 antibody was used to specifically detect the EWSR1-FLI1 protein expression, as FLI1 is not normally expressed in EWS cells (note that this is a common strategy in EWS research) (Garcia-Dominguez *et al.*, 2018; Smith *et al.*, 2006). We observed that ACY-1215 treatment reduced EWSR1-FLI1 expression levels in tumor samples, both as a monotherapy or in combination with doxorubicin (Fig. 4, E and F). While the reduction of EWSR1-FLI1 expression was higher in the combination therapy than in the ACY-1215 monotherapy, this difference was not statistically significant (Fig. 4E).

Collectively, these findings indicated that HDAC6 inhibition delays tumor growth *in vivo* and suggest that the anti-tumor activity was mediated, at least in part, through its inhibitory effects on the EWSR1-FLI1 oncogene expression.

## DISCUSSION

Due to the intrinsic characteristics of EWS, restoring epigenetic changes and inhibiting the oncogenic fusion protein could be a promising therapeutic alternative for EWS patients (Gierisch *et al.*, 2019; Herrero-Martin *et al.*, 2009; Kovar *et al*, 2016; Martinez-Lage *et al.*, 2020; Nacev *et al*, 2020; Scotlandi *et al*, 2020; Van Mater & Wagner, 2019). EWSR1-FLI1 enhances HDAC activity and inhibits HATs, thereby reducing histone acetylation levels (Sakimura *et al.*, 2005). Additionally, acetylation of EWSR1-FLI1 regulates the DNA binding capacity of the chimeric transcription factor (Schlottmann *et al*, 2012). These findings highlight an important role of HDACs in EWS and could contribute to the high sensitivity of EWS tumor cells to HDAC inhibitors.

In this study, we explored the use of HDAC inhibitors, based on the relevance of HDACs in cancer cell homeostasis (Parbin *et al*, 2014; Sun *et al.*, 2018). In fact, the use of pan-HDAC inhibitors in EWS cells induces EWSR1-FLI1 depletion and tumor growth inhibition (Garcia-Dominguez *et al.*, 2018; Sakimura *et al.*, 2005). However, this carries a high toxicity due to their effects on nonspecific targets (Hontecillas-Prieto *et al.*, 2020). Therefore, we hypothesized that it would be more appropriate to develop and evaluate the effects of selective HDAC inhibitors, which could maintain the effectiveness on EWSR1-FLI1 depletion while at the same time reducing treatment toxicity (de Nigris *et al*, 2020; Hontecillas-Prieto *et al.*, 2020; Sun *et al.*, 2018; Tang *et al*, 2017). We thus performed a screen of 43 epigenetic drugs, most of them known to modulate HDAC activity. Our results revealed that EWS cell lines were highly sensitive to BML-281, a specific HDAC6 inhibitor.

HDAC6 is emerging as a relevant HDAC in cancer, controlling several processes associated with oncogenesis, tumor maintenance, and regulation of autophagy/apoptosis (Mak *et al*, 2012; Sharif *et al*, 2019). We found that EWS cell lines were more sensitive to HDAC6 selective inhibition than other malignant cell lines and the non-tumor cell line hMSC (which are defined as the putative cells of origin for EWS (Tirode *et al*, 2007)). This emphasizes the importance of HDAC6 in EWS malignancy. Further, we revealed that the sensitivity of EWS cells to HDAC6 selective inhibition was associated with the presence of EWSR1-FLI1 in the ectopic HeLa model (Garcia-Dominguez *et al.*, 2020) after BML-281 treatment, but not with the levels of HDAC6 expression. Thus, the acquisition of this high sensitivity could be explained by the chimeric protein redrawing the epigenetic landscape, and thereby modulating the specific binding sites of HDACs (Pattenden *et al*, 2016). The specific HDAC targets regulated by EWSR1-FLI1 activity provide advantages to EWS tumor cells, and blockade of this regulation via HDAC inhibition could specifically sensitize EWS cells to BML-281 treatment.

Even though HDAC6 activity has been described mainly in the cytoplasm (Lee *et al.*, 2008; Seidel *et al.*, 2015), there is increasing evidence about its nuclear location (Li *et al*, 2008; Mobley *et al*, 2017; Wang *et al.*, 2009; Yang *et al.*, 2015). Here, we demonstrated the nuclear presence of HDAC6 in EWS cell lines by cell fractionation. In addition to the inhibition of HDAC6 activity in the cytoplasm, as assessed by α-tubulin acetylation, we found that BML-281 induced nuclear inhibition of HDAC6 activity and led to a significant increase of H4K12 acetylation. Given that nuclear activity of HDAC6 in EWS cell lines, as well as the EWS sensitivity to HDAC6 inhibition were associated with the presence of EWSR1-FLI1, we explored a possible role of HDAC6 in the regulation of EWSR1-FLI1 expression. We observed that BML-281 treatment significantly downregulated EWSR1-FLI1 mRNA level at 12 hours, and its protein level, at 24 hours, after treatment. In line with EWSR1-FLI1 downregulation, fusion-induced and fusion-repressed targets were down- and upregulated, respectively. Additionally, the expression signature induced by BML-281 in EWS cell lines as compared to the GSEA gene sets (C2_MSigDB) revealed a significant correlation with several previously published EWSR1-FLI1 target gene signatures. This response would explain why the use of pan-HDACis, such as SAHA or FK228, inhibits the driver of EWS (Garcia-Dominguez *et al.*, 2018; Sakimura *et al.*, 2005)—namely, due to the inhibition of HDAC6 activity by these drugs, as previously described (Furumai *et al*, 2002; Negmeldin *et al*, 2017).

The next step was to understand how HDAC6 regulates EWSR1-FLI1 expression. The *Let-7* genes modulation could likely contributes to *EWSR1-FLI1* regulation as a late effect of HDAC6 inhibition. However, another mechanism would be the main regulator of chimeric protein under the action of HDAC6 inhibition, both after short- and long-term treatment times, as previously observed. Indeed, we showed that the endogenous (non-translocated) *EWSR1* gene was also downregulated by HDAC6 inhibition, similar to inhibition of the fusion gene. Conversely, BML-281 treatment induced overexpression of the fusion protein in HeLa model. This excludes that post-transcriptional regulation of the fusion gene mRNA is a mechanism of *EWSR1-FLI1* depletion via HDAC6 inhibition; rather, it points to the *EWSR1* promoter as a possible HDAC6 target (as it is the differentiating element in this ectopic model with respect to the endogenous *EWSR1-FLI1* gene in EWS cell lines). Overall, these results suggest that regulation of the fusion gene by HDAC6 occurred via the *EWSR1* promoter (the homologous sequence in both endogenous and fusion genes in EWS). We verified this relationship using a gene reporter assay showing a dose-dependent luciferase gene inhibition (under the *EWSR1* promoter) by BML-281 treatment.

As HDAC inhibition induces transcription activation by restoring the acetylation levels of histones and non-histone proteins (Sun *et al.*, 2018), an intermediate element between HDAC6 activity and *EWSR1* promoter inhibition is necessary. Recent studies have demonstrated that SP1, a relevant transcription factor in cancer (Beishline & Azizkhan-Clifford, 2015), regulates *EWSR1-FLI1* (Giorgi *et al.*, 2015). Furthermore, Giogi et al. (2015) showed that SP1 depletion induces downregulation of the fusion gene and suggested that it is regulated via the fusion gene promoter (Giorgi *et al.*, 2015). Here, we demonstrated using chromatin immunoprecipitation assays that HDAC6 modulates binding of SP1 to the *EWSR1*/*EWSR1-FLI1* promoters. As a transcription factor, SP1 is able to recruit other elements, such as p300, to activate gene transcription (Azahri *et al.*, 2012; Formisano *et al.*, 2015; Hung *et al.*, 2006). In the same way, p300 binds to the *EWSR1* promoter in the absence of HDAC6 inhibitor (inducing *EWSR1*/fusion gene transactivation). Inhibition of HDAC6 releases both factors, downregulating both the fusion gene and the endogenous *EWSR1* expression. The loss of SP1 affinity to the *EWSR1* promoter by HDAC6 inhibition might be due to the acetylation of the only known residue, lysine 703, which resides in its DNA binding domain (Waby *et al.*, 2010). Waby et al. (2010) demonstrated that SP1 acetylation at lysine-703 releases this transcription factor from specific promoter targets (Waby *et al.*, 2010). Indeed, we observed increased SP1 lysine acetylation after BML-281 treatment. We suggest that HDAC6 inhibition induces the acetylation of the SP1 DNA-binding domain. This induction would then promote the release of SP1 and the recruited p300 from the *EWSR1*/*EWSR1-FLI1* promoters. Our results have shown for the first time a clear and specific epigenetic mechanism that regulates *EWSR1-FLI1* expression at its promoter, with HDAC6 playing a central role, and confirm the importance of using a selective HDAC6 inhibitor in EWS. This is in accordance with recent epigenetic marks that have been described to activate the fusion gene transcription: enrichment of H3K4me3, H3K9ac, and H3K27ac, which can result from SP1/P300 activity at the fusion gene promoter (Montoya *et al*, 2020). Whether the histone modification H4K12ac, a specific HDAC6 target, plays a role in the *EWSR1-FLI1* regulation remains to be investigated.

To determine the clinical relevance of HDAC6 in EWS, we evaluated the prognostic role of HDAC6 expression in patients with EWS. We demonstrated that HDAC6 overexpression correlated with poor outcome in these patients. Accordingly, we suggest the possible use of HDAC6 inhibition as an alternative treatment and an opportunity for EWS clinical trials. Using ACY-1215 treatment (a selective HDAC6 inhibitor that has successfully passed a phase Ib clinical trial for refractory multiple myeloma (Yee *et al*, 2016)), we observed a significant tumor growth inhibition in xenograft models. We used the A673 cell line and the EWS PDX derived primary cells, HSJD-ES-001, as the most representative models of EWS disease. The results obtained were positive using each xenograft model. HDAC6 selective inhibitors are known to affect protein trafficking, degradation, cell morphology, and migration, all of which are essential in tumor progression (Seidel *et al.*, 2015). Alternatively, their effect in EWS could also be explained by the fusion protein modulation via inhibition of HDAC6 activity, leading to alteration of HDAC aberrant chromatin accessibility regulated by the fusion protein in EWS. (Pattenden *et al.*, 2016) Thus, HDAC6 inhibition could be used as a selective therapy against the EWSR1-FLI1 epigenetic signature (de Nigris *et al.*, 2020).

To increase the efficiency in preclinical and clinical studies, HDAC6 selective inhibitors have been combined with DNA damaging or immunomodulatory drugs in several malignancies (Huang *et al*, 2017; Lee *et al*, 2018; Ray *et al*, 2018). Here, we combined ACY-1215 with doxorubicin (a DNA damage–inducing agent), which is a standard drug for EWS patient treatment. This combination enhanced tumor growth inhibition as compared to either monotherapy. Moreover, HDAC6 inhibition *in vivo* induced the EWSR1-FLI1 protein downregulation under ACY-1215 treatment, both in monotherapy and in combination. These results validated the *in vitro* results, leading us to postulate that HDAC6 is a EWSR1-FLI1 inhibitor *in vivo*. Additionally, based on general behavior, weight, and organ histopathology evaluation, no toxicity was present in our experimental mice with either ACY-1215 alone or in combination with doxorubicin. Indeed, HDAC6-deficient mice are viable and fertile (Zhang *et al.*, 2008). We therefore believe that HDAC6 depletion can be a safe strategy for EWS patient treatment.

In light of all our results, we consider that selective HDAC6 inhibition focused on restoring aberrant epigenetic changes and on inhibition of the oncogenic EWSR1-FLI1 fusion transcription factor could be a promising, selective, and safe EWS therapeutic alternative.

## MATERIALS AND METHODS

### Epigenetic drug library screening

A4573, CADO-ES, RDES, SK-ES-1, SKNMC, STAET 2.1, and TTC466 EWS cells were seeded in 96-well plates and treated with 43 different epigenetic drugs. Non-treated cells were resuspended in medium with DMSO (0.1%). Proliferation was measured at 72 hours after treatment by MTT assay, and absorbance rates at 595 nm were measured by spectrophotometer (BIO-RADiMark). This screening was conducted by the Mejoran Lab (Madrid).

### Acid histone extraction

Cells were washed with ice-cold 5 mM sodium butyrate-PBS (1×). Centrifuged and resuspend cells in hypotonic extraction buffer (10 mM Tris-Cl pH 8.0, 1 mM KCl, 1.5 mM MgCl_2_, and 1 mM DTT) were supplemented with protease inhibitor. Cells were lysed on ice for 30 min and centrifuged for 10 min at 10,000*g* for nucleus precipitation. Supernatants were discarded and eluted in 0.4 M H_2_SO_4_, incubated for 30 min on ice, and then centrifuged 10 min at 16000*g*. Supernatants were treated with 33% trichloroacetic acid (v/v) and incubated overnight at –20°C to precipitate histones. Pellets containing histones were washed twice with cold acetone and then centrifuged 5 min at 16000*g*, and supernatants were removed and air dried. Histone pellets were resuspended in ddH_2_O.

### Short hairpin RNA (shRNA) HDAC6

Two pLKO.1-shRNA constructions against the HDAC6 transcript were selected from the MISSION shRNA collection (SIGMA Aldrich). The constructs references were TRCN0000314976 for sh1_HDAC6 and TRCN0000004839 for sh2_HDAC6. An empty MISSION pLKO.1-puro (SIGMA Aldrich) was included as a control of the off-target effects. These pLKO.1 constructions were transfected in the 293T packaging cell line along with pMD2.G and pCMV-dR8.91 vectors (Addgene), following a typical Lipofectamine 2000 (Invitrogen) protocol. At 48 hours after transfection, viral supernatants were collected and fresh medium was added; they were then filtered through a 0.44-μm polysulfonate filter (PALL), with polybrene added to a final 8 μg/ml concentration (SIGMA Aldrich). These filtrates were used to transduce the target EWS cell lines (4 × 10^5^ cells seeded per well in 6-well plates). Plates were centrifuged at 400*g* for 1 hour at 32°C. After 72 hours, transfected cells were selected by puromycin addition to medium.

### Next-generation transcriptome sequencing (RNA-seq) for gene expression analysis

Total RNA from 18 samples were sequenced in a NextSeq 500 sequencer (Illumina, CA, USA) producing 23,830,047 raw 75 × 2 nt paired-end reads on average. Sequences were processed using miARma-Seq pipeline (Andres-Leon *et al*, 2016). Briefly, quality filtered reads were aligned using HISAT2 (Kim *et al*, 2015), resulting in a 90.13% of properly aligned reads that were summarize into gene expression values using featureCounts (Liao *et al*, 2014). Differential expression analysis was done using the edgeR package (Nikolayeva & Robinson, 2014), generating differentially expressed genes (DEG) between time points, using both tissues at 4 hours and 24 hours (6 replicates per time) versus control conditions (6 replicates). Next, a Gene Set Enrichment Analysis (GSEA) (Subramanian *et al*, 2005) was performed to understand the resulting expression profile considering DEG (FDR < 0.05 and logfc>|1.5|). Normalized enrichment scores NES were calculated between DEG against a curated dataset of MSigDB, called C2 CGP, which comprises 3358 gene sets representing expression signatures of Genetic and Chemical Perturbations (see supplementary information for further details).

### Luciferase reporter assay

For luciferase assays, the pGL3-Basic vector (Promega) was used, and the *EWSR1*-activating promoter region (–2000 bp to TSS (Moller *et al.*, 2009)) was successfully cloned upstream of the luciferase gene. pGL3-Empty vector was included as a control for off-target effects. Cells were seeded in 96-well plates at 24 hours before transfection. Both constructs were transfected with Lipofectamine 2000 (Invitrogen) together with pRL-TK Renilla (100 ng/well) to normalize for transfection efficiency. Cells were treated for 4 hours with different doses of BML-281 or control medium (DMSO 0.1%) and then collected at 4 hours after treatment. Cell lysates were assessed using the Dual-Luciferase Reporter Assay System (#E1910; Promega, Madison, WI, USA). Luminescence was measured using a TECAN M200 infinite-PRO plate reader (Tecan, Männedorf).

### Chromatin immunoprecipitation (ChIP)

EWS cells were fixed with 1% formaldehyde at room temperature for 10 min, followed by 5 min quenching with 0.125 M glycine, and then washed once in ice-cold PBS. Pellets were resuspended in lysis buffer (0.1% SDS, 0.1 M NaCl, 1% Triton X-100, 1 mM EDTA, 20 mM Tris pH 8, and 1 mg/ml protease inhibitors) and sonicated with a Bioruptor until the crosslinked chromatin was sheared, with an average DNA fragment length of 0.5 kbp. After centrifugation (30 min at 15,700 *g*), chromatin preparations were precleared by incubation with 40 μl of protein A agarose/salmon sperm DNA, 50% gel slurry (Millipore) for 2 hours at 4°C under rotation. Protein A agarose was removed by centrifugation, and the precleared chromatin was immunoprecipitated by incubation with 5 μg anti-SP1 or anti-p300 antibodies and 50 μl protein A agarose overnight at 4°C. Washed pellets were eluted with 120 μl of a solution containing 1% SDS, 0.1 M NaHCO_3_. Eluted pellets were de-crosslinked at 65°C overnight and purified in 50 μl Tris-EDTA buffer using the QIAquick PCR Purification Kit (Qiagen). Differences in the DNA content from every immunoprecipitation assay were determined by real-time PCR using the ABI 7700 sequence detection system and SYBR Green master mix protocol (Applied Biosystems). Primers used in this study are listed in table S1. The reported data represent real-time PCR values normalized to input DNA and expressed as percentage (%) of bound/input signals.

### Clinical samples and tissue microarrays (TMAs)

EWS samples were acquired from the Department of Pathology at the Hospital Universitario Virgen del Rocío (Seville, Spain) and the HUVR-IBiS Biobank (Seville, Spain). The series of EWS samples were obtained between 1991 and 2017 and comprised 52 paraffin-embedded tumor samples. Patient characteristics are summarized in table S2. Approval of the Ethics Committee of our institution was obtained, and written informed consent was obtained from each patient before including samples and data into the HUVR-IBiS Biobank.

Tissue sections (5 μm) from formalin-fixed paraffin-embedded (FFPE) of EWS tumors were stained with hematoxylin & eosin. Representative malignant areas from samples were carefully selected from the stained sections of each tumor, and two 1-mm diameter tissue cores were obtained from each sample, to build up TMAs in duplicate.

### Statistics

Differences between control and treatment conditions were evaluated using Mann–Whitney U-test for two groups, and one-way analysis of variance test for more than two groups followed by post hoc Tukey’s multiple comparisons. The disease-free survival (DFS) time was analyzed using the Kaplan– Meier estimator and the Wilcoxon test. For all analyses, *P* values of ≤ 0.05 were considered statistically significant. Analyses were performed using the Prism 6 software (GraphPad). All experiments were performed in triplicate, with each repeated at least three times.

Additional methods are described in Supplementary Material and Methods.

## ACKNOWLEDGMENTS

The authors thank the donors and the HUVR-IBiS Biobank (Andalusian Public Health System Biobank and ISCIII-Plataforma de Biobancos PT13/0010/0056) for the human specimens used in this study.

## AUTHOR CONTRIBUTIONS

D.G.D. and L.H.P. conceived the study and its design, analyzed the data, and wrote the manuscript. D.G.D., S.S.M., E.F.B., R.F.C., and L.H.P. performed most experimental work. D.G.D., R.M.P., A.M.E., G.P.P., A.M.C., and L.H.P. carried out the work with animals. E.A.L. and L.C.T. performed the bioinformatics analyses. M.J.R. and E.D.A. performed the anatomopathological evaluation. N.B., J.M., and E.D.A. participated in the conceptualization of the study and in the critical discussion of data. All authors revised and edited the manuscript.

## DATA AVAILABILITY STATEMENT

Our RNA-seq data have been deposited in the NCBI Sequence Read Archive (SRA) as PRJNA673347: (https://dataview.ncbi.nlm.nih.gov/object/PRJNA673347?reviewer=ktphm151lrr8mu1ol2jml0go7s).

## TABLES LEGENDS

**Table 1. Summary of RNA-seq results for BML-281–treated SKNMC and WE68 cell lines at 4 hours or 24 hours after treatment.**

**Table 2. Functional enrichment analysis based on gene ontology (GO) for BML-281–treated SKNMC and WE68 cell lines at 4 hours or 24 hours after treatment.**

**Table 3. GSEA of rank-ordered gene sets (C2_MSigDB) in BML-281–treated SKNMC or WE68 signatures at 24 hours after treatment.** Ranked gene set: NOM *P* < 0.05.

